# Considerations and complications of mapping small RNA libraries to transposable elements

**DOI:** 10.1101/079749

**Authors:** Alexandros Bousios, Brandon S. Gaut, Nikos Darzentas

## Abstract

The advent of high-throughput sequencing (HTS) has revolutionized the way in which epigenetic research is conducted. Often coupled with the availability of fully sequenced genomes, millions of small RNA (sRNA) reads are mapped to regions of interest and the results scrutinized for clues about epigenetic mechanisms. However, this approach requires careful consideration in regards to experimental design, especially when one investigates repetitive parts of genomes such as transposable elements (TEs), and especially when such genomes are large as is often the case in plants. Here, to shed light on the challenges of mapping sRNAs to TEs, we focus on the 2,300Mb maize genome, of which >85% is derived from TEs. We compare various methodological strategies that are commonly employed in TE studies. These include choices for the reference dataset, the normalization of multiple mapping sRNAs, and the selection among different types of sRNA metrics. We further examine how these choices influence the relationship between sRNAs and the critical feature of TE age, and explore and contrast their effect on low copy regions (exons) and other popular HTS data (RNA-seq). Finally, based on our analysis, we share a series of take-home messages to help guide TE epigenetic studies specifically, but our conclusions may also apply to any work that involves mapping and analysis of HTS data.

## INTRODUCTION

Across eukaryotes, epigenetic pathways contribute to diverse functions, including gene regulation and transposable element (TE) silencing (1). Small RNAs (sRNAs) are a key component of these pathways. Numerous studies have investigated the biogenesis and functional roles of sRNAs, with most focusing on the molecular mechanisms that underlie these processes (for recent reviews see (2–4)). Some of these studies have utilized high-throughput sequencing (HTS) technologies, which generate vast numbers of sRNA reads. This capacity of HTS has facilitated the identification of novel sRNA classes, the quantification and comparison of sRNA expression profiles across tissues, and the discovery of genomic loci that map large volumes of sRNAs. These tasks have been aided by numerous computational tools, most of which have been tailored to study micro RNAs (miRNAs) (5–11), with fewer offering comprehensive identification, quantification and visual-based support for all sRNA types (12–17).

Even with these tools, significant challenges remain in the handling and interpretation of HTS sRNA data. One major challenge concerns the sequencing depth of libraries during differential expression analysis across tissues; this topic, however, has been thoroughly investigated (e.g., (6,15,18) and will not be discussed further.

Another important challenge, which we will address here, stems from the fact that some sRNAs map to unique locations (U_sRNAs) of a reference genome, while others align equally well to multiple locations (M_sRNAs). The handling of the latter is a major concern, as it impacts downstream analyses (15), and is as yet practically unresolved as different studies (reviewed in Johnson et al. 2016) and related sRNA analysis tools have used different approaches. For example, the NiBLS method allows multiple mapping without any kind of normalization for the number of mapping locations (19), the SiLoCo tool of the UEA sRNA Toolkit weights each read by its repetitiveness in the genome (20), the segmentSeq package of Bioconductor allocates each M_sRNA only once to a predefined locus even if it maps to more than one places within this locus or indeed across the genome (13), Novoalign (http://www.novocraft.com) excludes M_sRNAs, and bowtie (21) and bwa (22) randomly place each M_sRNA to a single locus under their default settings (22). Finally, a recently updated version of ShortStack allocates M_sRNAs to single loci based on the densities of U_sRNAs (23).

The importance of M_sRNAs and of their handling may be dependent on the component of the genome under investigation; for instance, due to their repetitive nature, TEs are likely to map many M_sRNAs, which unavoidably complicates TE-related studies. This effect may be especially prominent in plants, because of their large genomes (the average size of a diploid angiosperm is ~6,400Mb) and the fact that most of plant DNA has originated from TEs (24). This point is exemplified by contrasting data from the unusually small genome of *Arabidopsis thaliana* (which is only 125Mb and contains ~24% TE-derived DNA) and the larger - but still small, relative to the angiosperm average - genome of maize (2,300MB, ~85%). sRNA mapping studies have shown that <25% of *A. thaliana* TEs are mapped solely by M_sRNAs (25), but this proportion raises to >72% for maize TEs (26). Hence, careful consideration of M_sRNAs is crucial for understanding epigenetic processes in genomes like maize. The challenges of mapping sRNAs to TEs are further exacerbated by the fact that accurate TE identification is a notoriously difficult task (27). To simplify the problem, previous studies have often used TE exemplars (28–30), which are a consensus of many TE sequences representing a single TE family or subfamily. The use of exemplars may be pragmatic, but it likely reduces the analysis resolution compared to examining whole populations of annotated TEs.

Here we attempt to systematically address the complex, but understudied, issue of analyzing sRNAs in the context of TEs, because the impact of their treatment on analyses is presently unclear. To better assess different approaches, we focus on the maize genome and the most abundant *Copia* and *Gypsy* Long Terminal Repeat (LTR) retrotransposon families. We perform standard sRNA mapping using existing HTS data from three different tissues, but vary several features of the analyses, such as i) the reference dataset, which ranges from whole genome TE annotations to TE exemplars, ii) the treatment of M_sRNAs, which ranges from various normalization options to their complete exclusion, and iii) the sRNA metrics, i.e. consideration of distinct sequences or their abundances. Figure 1 depicts the methodological matrix of our work, along with many of the terms that we use throughout the study. We then comment on the effect of some of these choices on the relationship of mapping with other TE features such as TE age, with low copy regions of the maize genome, or when using HTS mRNA data. We conclude by sharing our insights as take-home messages to help guide researchers in epigenetic analyses of TEs or mapping of HTS data to large and complex genomes.

**Figure 1.**
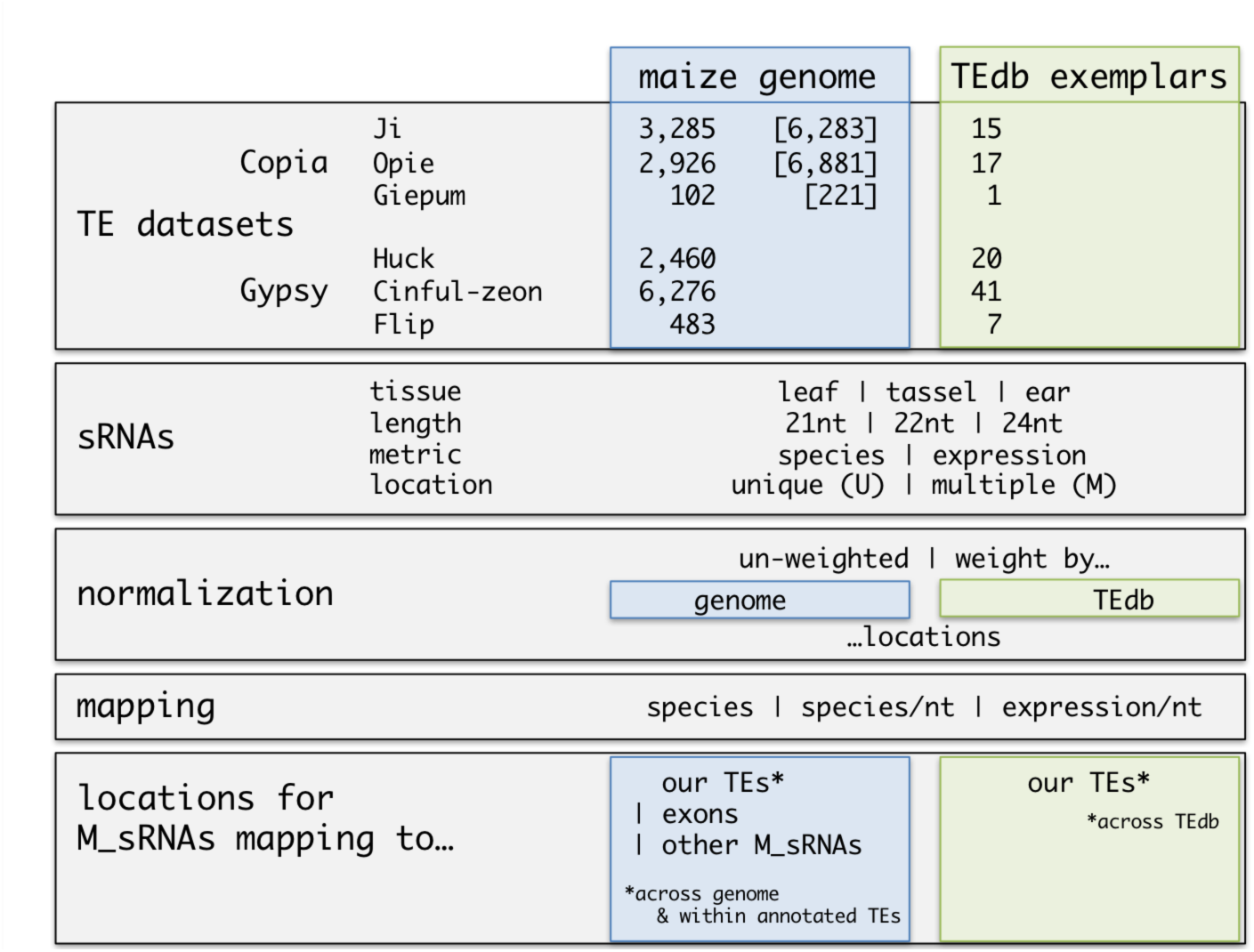
A matrix of the terms, data and analyses used in this study.

## RESULTS & DISCUSSION

### Reference datasets: TE exemplars vs. annotated TE populations

It is common practice in TE epigenetic studies to examine only full-length TEs. But how do inferences vary as a function of the reference dataset? In this study, we compared mapping patterns of 21, 22 and 24 nucleotide (nt) sRNAs of leaf, tassel and ear tissues between two ΤΕ datasets. The first was a comprehensively annotated dataset of Sirevirus LTR retrotransposons of the *Copia* superfamily in maize (31). Sireviruses encompass the three most abundant *Copia* families in maize, namely *Ji*, *Opie* and *Giepum*. *Ji* and *Opie* each constitute ~10% of the genome, and *Giepum* represents another ~1.2% (32,33). We used 3,285 *Ji*, 2,926 *Opie* and 102 *Giepum* full-length elements that were recently analyzed for their epigenetic patterns (26) (Figure 1). To add to this dataset, we devised a pipeline (see Methods) to identify full-length elements of the three most abundant *Gypsy* families, including *Huck* (10.1% of the genome), *Cinful-zeon* (8.2%) and *Flip* (4.2%) (32) (Figure 1). We annotated 2,460, 6,276 and 483 copies respectively. The second reference dataset consisted of exemplar TE sequences. We downloaded all maize TE exemplars from http://maizetedb.org. The number of exemplars for the six *Copia* and *Gypsy* families ranged from one to 41 consensus sequences (Figure 1).

### Exemplars miss a substantial fraction of sRNA information

We began by first examining the total number of distinct sRNA sequences (termed “sRNA species” hereafter) mapped to each family. An initial observation was that there is a much lower number of sRNAs (3-fold decrease on average) that mapped to the exemplars compared to the annotated populations (Figure 2A, Table S1). For example, 90,503 sRNA species of the leaf library mapped to the exemplars of all six families combined, compared to 310,548 that mapped to the annotated elements. This result indicates that a substantial fraction of sRNA information is lost when using exemplars. Nonetheless, the majority of sRNAs that mapped to exemplars do relate to the six families, because 89% (162,493 of 181,188 for all libraries and families combined) also mapped to the annotated TE populations.

**Figure 2.**
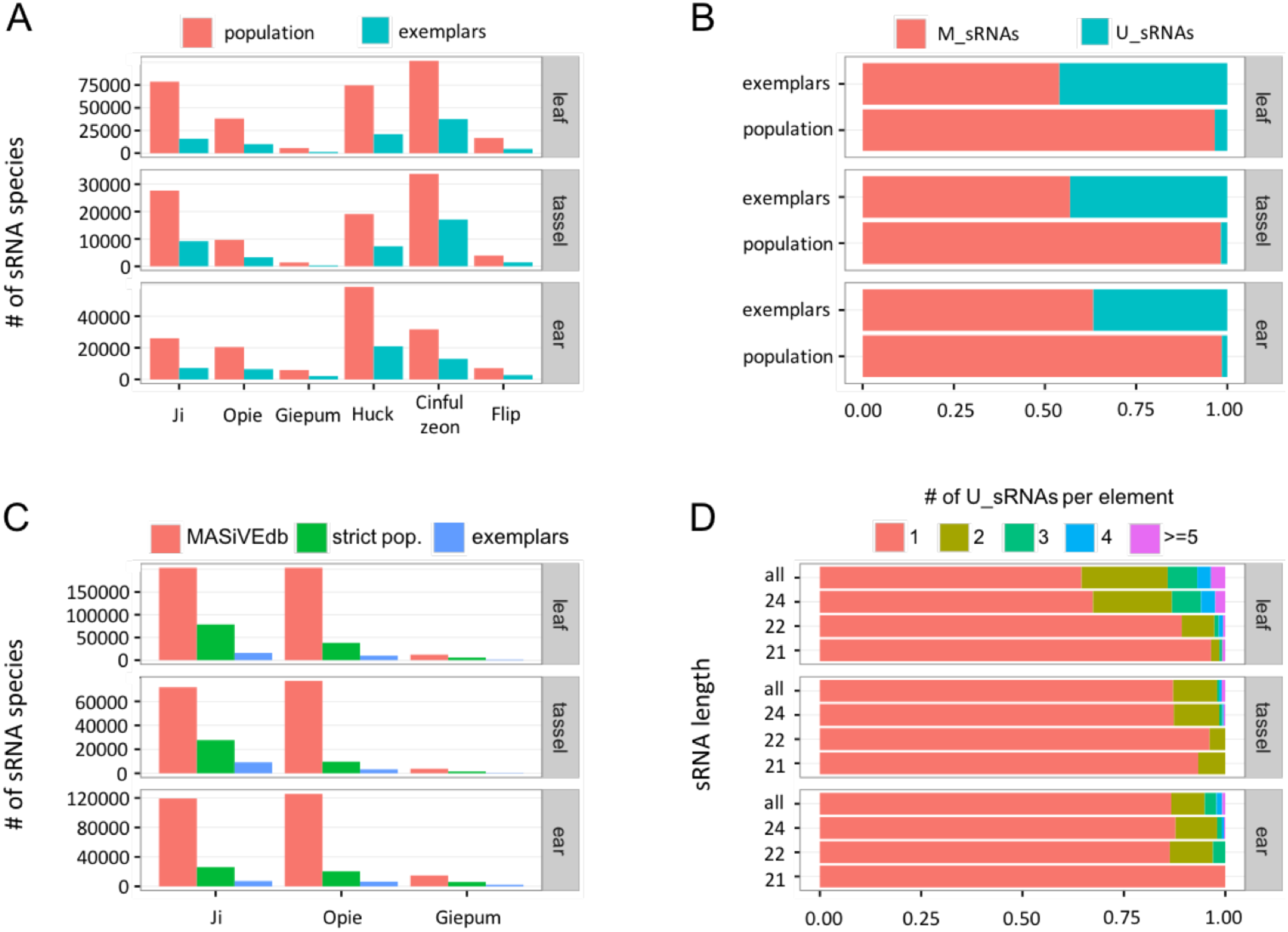
sRNA metrics on TE exemplars and annotated TE populations. (**A**) Total number of sRNA species that mapped to each family. (**B**) Proportion of U_sRNAs and M_sRNAs for all families combined. (**C**) Total number of sRNA species that mapped to different datasets of the three *Copia* families. (**D**) Proportion of the number of U_sRNAs that mapped per TE.

### U_sRNA and M_sRNA ratios differ between datasets

Previous research has suggested that U_sRNAs may exert a stronger effect on TE silencing compared to M_sRNAs (25,34). Accordingly, several studies have used only U_sRNAs as the basis for inference, derived either from mapping to genomes or to exemplars (29,30,35,36). Our analysis showed that there is a massive difference in the U:M sRNA ratio as a function of the reference dataset: a much higher proportion of sRNAs map uniquely to exemplars (43% of all sRNAs for all libraries and families combined) compared to annotated TE populations (2.6%) (Figure 2B, Table S2). In fact, the vast majority of U_sRNAs that map to exemplars become M_sRNAs when mapped to the genome. This finding raises two important points. First, U_sRNAs appear to be overrepresented in the analysis of exemplar datasets and hence their use over M_sRNAs (e.g., (29,30)) should be carefully considered. Second, U_sRNAs may convey little information at the genome level for TE studies, because there may be only a few mapping to each element, at least in TE-rich and large genomes like that of maize.

### Low levels of sRNA ‘cross-talk’ between families are consistent between datasets

The extent to which sRNAs can map to elements of multiple TE families is largely unknown. Yet, this epigenetic ‘cross-talk’ may be important for host defenses, by spreading TE silencing through homology-based mechanisms (37,38). Our analysis of both TE exemplars and annotated TE populations was consistent in showing that sRNAs are generally family specific. For all libraries combined, only 0.2% of 181,188 exemplar-mapping and 1.7% of 550,755 population-mapping sRNA species cross-talked between families. For completeness, we also investigated the degree of sRNA cross-talk between TEs and exons from the maize Filtered Gene Set. The low levels of cross-mapping (<0.2% for each library) imply that exon-mapping sRNAs (545,601 sRNAs for all libraries combined) serve biological roles unrelated to TE silencing.

### sRNA patterns along TE sequences differ between datasets

We then examined the mapping characteristics of each TE dataset by averaging sRNA targeting per nucleotide of each full-length element, so as to allow comparisons among TEs. For this analysis, we used sRNA species but also the metric of “sRNA expression” that indicates the number of reads in a library for each sRNA species. In addition, we did not normalize M_sRNAs by their number of mapping locations. (We compare weighted and un-weighted data further below). Our analysis showed that, despite the low number of sRNAs that mapped to exemplars, there was agreement across datasets (Figure S1): 24nt sRNAs, measured either as species or expression, generally targeted all families more intensely than 21-22nt sRNAs, although this was not as evident in tassel and for *Ji* that was previously categorized as a “22nt” family (39).

Next, we examined if this consistency persists during in-depth analysis of sRNA mapping along the length of elements. For this comparison we focused on the three *Copia* families, because of the preexisting annotation of their genomes, including information about complex palindrome motifs in the *cis*-regulatory region of the LTRs that are sRNA targeting hotspots (26). We found that both datasets produced highly similar patterns, with one intriguing exception: the exemplars were not mapped by sRNAs in the palindrome-rich regions (Figure 3A). Closer investigation of the exemplar sequences revealed that they contain long runs of N nucleotides in these regions (Figure 3B). We presume that the process of generating consensus sequences does not perform well in areas of high sequence variability, because the palindromes of the Sirevirus families differ both along the sequence of an individual element and among elements (26). As a result, these regions are masked in exemplars, even though they may be of biological importance due to their elevated sRNA mapping and rapid evolution (26).

**Figure 3.**
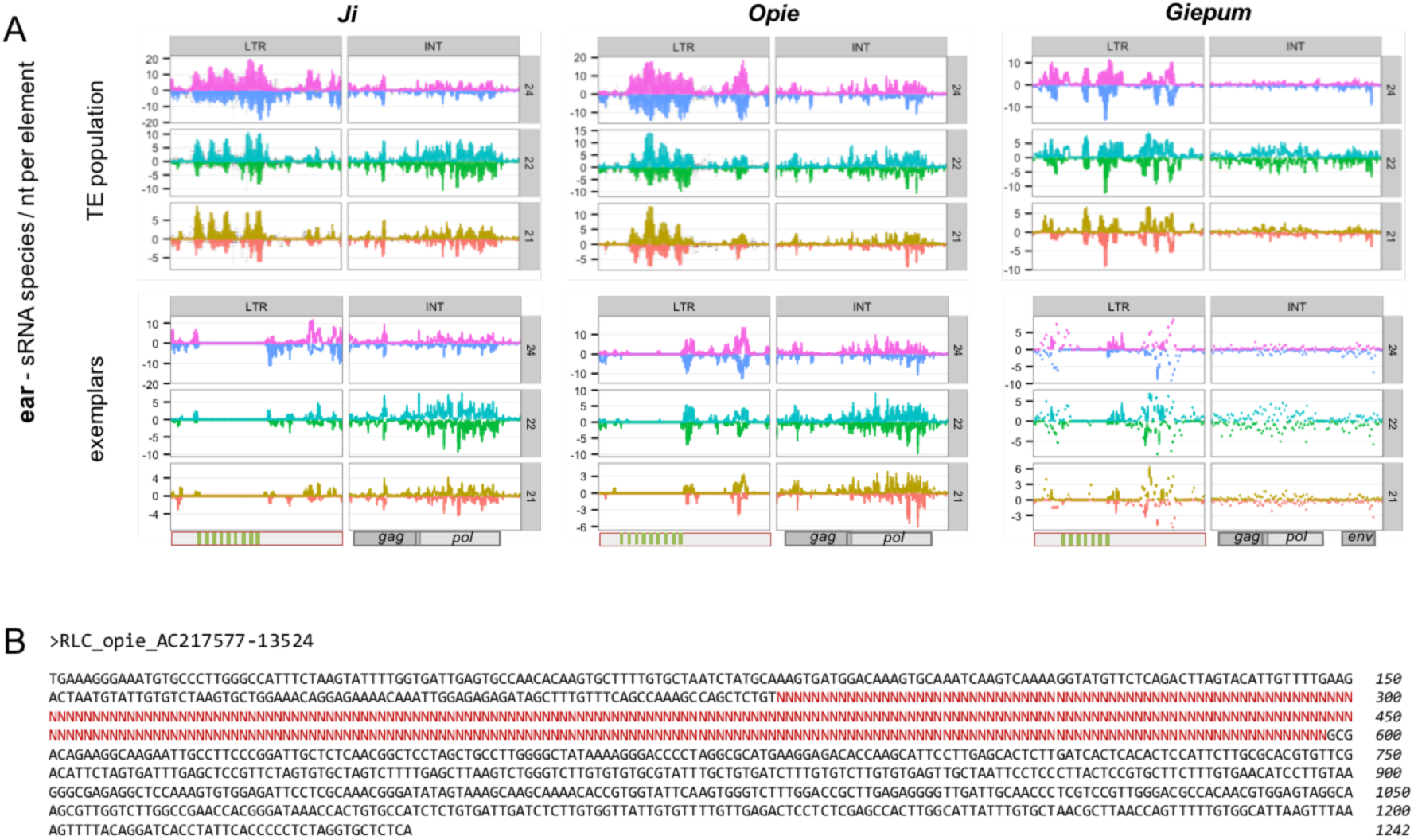
sRNA mapping along the sequences of *Ji*, *Opie* and *Giepum* exemplars and annotated populations. (**A**) Un-weighted sRNA data from ear tissue were mapped separately to the LTRs and the internal (INT) domain. Each region was first split in 100 equally sized windows, and mapping was calculated as the number of sRNA species per nucleotide of the sense (positive *y*-axis) and antisense (negative *y*-axis) strands, and visualized with a boxplot for each window. The position of the palindromes (LTRs) and the *gag, pol* and envelope (*env*) genes (INT domain) are shown at the bottom of each panel. (**B**) An example of the LTR sequence of an *Opie* exemplar with N nucleotides masking the unresolved palindrome-rich region.

Searching further, we found that 74 exemplars from 37 families within http://maizetedb.org contain stretches of >100 N nucleotides (*Huck*, *Cinful-zeon* and *Flip* were not among them), making the occurrence of masked regions a fairly common feature of this dataset. The extent of this problem is not known for other plant species that have generated exemplar datasets such as foxtail millet (40) and strawberry (41), but needs to be assessed, especially in the light of how helpful these datasets can be in combination with genomic, sRNA and mRNA HTS data in the analysis of the repetitive fraction of genomes (42,43). Overall, this analysis illustrates that the use of exemplars may not only reduce the volume of available sRNA information (Figure 2A, Table S1), but may in fact entirely omit mapping to specific regions of TEs (Figure 3A).

### ‘Contamination’ of annotated TE populations affects sRNA mapping profiles

Our reference dataset representing the three *Copia* families is a strictly curated subset of the complete population of maize Sireviruses available from MASiVEdb (http://bat.infspire.org/databases/masivedb/). This larger dataset contains 6,283 *Ji*, 6,881 *Opie* and 221 *Giepum* full-length elements (Figure 1) that have been identified as *bona fide* Sireviruses (44). However, unlike our reference dataset, these TEs have not been carefully examined for the presence of ‘contaminating’ DNA from insertions of other elements or even from captured genes. As a result, an unknown number of elements may harbor foreign DNA, which in turn may affect data interpretation.

To examine this issue, we compared the mapping characteristics of the reference dataset against the complete MASiVEdb population. The number of sRNA species that mapped to each TE family increased substantially for MASiVEdb. Collectively, 626,836 sRNAs from the three sRNA libraries mapped to the 13,385 TEs of MASiVEdb, but only a third (206,589) of that total mapped to our reference dataset (Figure 2C, Table S1). Hence, the MASiVEdb population represents far more sRNA variation, but the extent to which TE contamination contributes to this increase is unclear. It is also hard to assess, because even very small fragments may map several sRNAs that could collectively account for a large fraction of the sRNA pool.

One indication may be provided by comparing the level of sRNA cross-talk between families of the two datasets. Our conjecture is that higher levels of cross-talk in MASiVEdb will reflect the presence of fragments of one family within elements of another family, thereby artificially increasing their ‘common’ sRNAs. Our analysis showed that indeed this was the case. For example, 30.8% (188,926 of 612,225 for all libraries combined) of the *Ji* and *Opie* sRNAs mapped to elements of both families using MASiVEdb compared to 3.1% (6,033 of 194,582) using the reference dataset. Likewise, cross-talk increased with the *Gypsy* families, for example from 0.2% to 5.3% between *Ji* and *Huck*, and from 0.2% to 10% between *Opie* and *Cinful-zeon*. To further test this conjecture, we screened for foreign TE fragments within the two datasets using non-Sirevirus maize TE exemplars as queries (BLASTN, max *E*-value 1×10^−20^). We detected only two elements of the reference dataset with foreign TEs, compared to 1,158 elements of MASiVEdb that contained fragments (of 189nt median length) from 451 non-Sirevirus families.

Using MASiVEdb as an example, this analysis implies that a substantial fraction of the full-length elements of TE databases contain foreign TE fragments. Yet, that is not to say that these databases are of low quality; they all employ sophisticated algorithms based on various structural and homology criteria to identify high-quality full-length elements. To support this argument, we note that MASiVEdb generated similar sRNA mapping patterns as the reference dataset along the TE sequence, including the palindrome-rich region (Figure S2). It is likely, however, that other types of epigenetic analyses might be negatively affected by TE contamination. Hence, it is advisable that the sequence quality of TEs is considered prior to mapping of sRNA data.

### Normalization: Complexities regarding the use of M_sRNAs

#### Inclusion of M_sRNAs is necessary in TE studies

The handling of sRNAs with multiple mapping locations is an issue that has long troubled scientists. Often, in an effort to avoid methodological complications, M_sRNAs are excluded from analyses (29,30,35,36). However, even though U_sRNAs correlate more consistently with TE silencing than M_sRNAs (25), a significant proportion of RNA-directed DNA methylation (RdDM) is thought to be mediated by M_sRNAs (34). Moreover, our data in Figure 2B suggest that there may not be enough U_sRNAs to make meaningful inferences about TEs in hosts with large genomes.

To examine potential U_sRNA differences among plant species, we calculated the median density of 24nt U_sRNAs per nucleotide of maize TEs (for all libraries and families combined) and compared it to those of *Arabidopsis thaliana* and *lyrata* TEs previously reported by Hollister et al. (2011). While the median densities were only twofold different between *thaliana* and *lyrata* (0.11 vs. 0.06), these two species had a 69-fold and 37-fold difference with maize respectively (0.0016 24nt U_sRNAs per nucleotide of maize TEs). Comparative data were not available for 21-22nt U_sRNAs from the Hollister et al. study, but given that only 3,522 21-22nt U_sRNAs from all libraries mapped to the 15,532 full-length elements of the *Copia* and *Gypsy* datasets combined, it is clear that most elements did not map U_sRNAs in maize. These findings i) suggest that the effect of U_sRNAs on TE silencing may be less prominent in maize and other large genomes, and ii) decisively argue in favor of including M_sRNAs in TE analyses.

#### Normalization of M_sRNAs varies across genomic regions and between datasets

Besides excluding M_sRNAs from analyses as discussed previously or sometimes even allocating them randomly to single loci (45–47), the most common approaches for handling M_sRNAs is either to count all mapping locations so that each location has a value of 1.0, or to weight for multiple mapping so that each location is assigned a value of 1/*x*, where *x* is the total number of locations for a given M_sRNA. This normalization can be applied to both “sRNA species” and “sRNA expression”. Nonetheless, it is unclear if and how these normalization strategies affect downstream research. One parameter that may provide valuable insights is the extent of multiple mapping (i.e., the number of mapping locations) for M_sRNAs that target various parts of a genome or different reference datasets (Figure 1). The reasoning is that the smaller the *x*, the weaker the differences between strategies will be and *vice versa*. To our knowledge, such a systematic examination has not been conducted so far. We therefore compared the mapping locations of Μ_sRNAs that target our *Copia* and *Gypsy* families i) across the genome, ii) within their annotated full-length populations, and iii) across the TE exemplar database, so as to keep in line with the various strategies of previous studies.

Focusing first on the entire genome, we find that Μ_sRNAs have an exceptionally high number of mapping locations. For example, the median number of locations for all families combined was up to 513 among the three libraries, while the average often exceeded 1,500 (Table 1). Interestingly, separate investigation of the *Copia* and *Gypsy* families revealed that Μ_sRNAs targeting *Copia* elements had twice as many mapping locations compared to *Gypsy* elements (Table S3), even though each group of families comprises ~20-25% of the maize genome (32,33). This discrepancy may reflect different levels of sequence heterogeneity, i.e. families with many highly similar elements provide substantially more mapping loci for a given Μ_sRNA, compared to families with more divergent elements. Second, in contrast to the numerous mapping locations across the entire genome, there was a marked decrease in the number of locations within the annotated full-length populations (Table 1). We found that, on average, only a fifth of the genomic locations correspond to full-length elements, indicating that most Μ_sRNAs map to other types of sequences related to the six families, presumably unidentified full-length elements, degraded copies or solo LTRs. Third, the decrease was even more dramatic within the TE exemplar dataset, where the M_sRNAs of the six families only had three to five mapping locations each (Table 1).

**Table 1.** Number of locations for M_sRNAs that mapped to different parts of the maize genome.

| library | sRNA length | # of loci for sRNAs of the six families<sup>a</sup> |  |  | # of genomic loci for exon sRNAs<sup>a</sup> | # of genomic loci for other sRNAs<sup>a</sup> |
| --- | --- | --- | --- | --- | --- | --- |
|  |  | genome | TE population | exemplar db |  |  |
| leaf | 21 | 283 - 1,397 | 66 - 298 | 3 - 5 | 4 - 12 | 5 - 37 |
|  | 22 | 262 - 1,261 | 70 - 284 | 3 - 5 | 4 - 11 | 5 - 42 |
|  | 24 | 82 - 613 | 11 - 121 | 3 - 4 | 4 - 12 | 4 - 21 |
|  | all | 127 - 854 | 18 - 179 | 3 - 4 | 4 - 11 | 4 - 26 |
| tassel | 21 | 425 - 2,033 | 114 - 419 | 3 - 5 | 4 - 18 | 6 - 57 |
|  | 22 | 380 - 1,615 | 118 - 369 | 3 - 5 | 4 - 15 | 7 - 60 |
|  | 24 | 199 - 1,017 | 26 - 194 | 3 - 4 | 5 - 17 | 4 - 25 |
|  | all | 277 - 1,353 | 60 - 281 | 3 - 5 | 5 - 17 | 4 - 34 |
| ear | 21 | 513 - 2,130 | 86 - 411 | 4 - 5 | 4 - 14 | 6 - 55 |
|  | 22 | 454 - 1,748 | 83 - 359 | 4 - 5 | 4 - 15 | 7 - 56 |
|  | 24 | 147 - 897 | 19 - 170 | 3 - 5 | 4 - 17 | 5 - 26 |
|  | all | 219 - 1,231 | 31 - 241 | 3 - 5 | 4 - 16 | 5 - 32 |

<sup>a</sup>The median (left) and average (right) number of mapping locations are shown for each category.

The above findings were derived from the most abundant TE families in maize and hence represent the most repetitive parts of a large genome. To contrast them with lower copy regions, we calculated the genomic locations of two additional sets of M_sRNAs: exon-mapping M_sRNAs and all other M_sRNAs that did not map to either exons or the six TE families. We assume that a substantial proportion of the last category corresponds to less abundant TE families. (Note that the libraries were filtered for tRNAs, rRNAs, snoRNAs and miRNAs). Our analysis showed that the mapping locations of both categories did not exceed a handful of sites (Table 1); nonetheless, the average number of locations of the ‘other’ M_sRNAs was three-fold higher than the exon-mapping M_sRNAs, implying that a large proportion of the former type may indeed target low copy TEs.

Altogether, these data show that there can be up to a hundred-fold variation in the number of locations for M_sRNAs of maize TEs, depending on the reference dataset that is employed. This huge variation particularly affects the most abundant TE families, because mapping to TE exemplar datasets turns extreme multi-mappers to essentially semi-U_sRNAs (Table 1). Moreover, this effect likely holds true for the majority of plants, as most have genomes larger than maize and with concomitant TE content (24). Based on this evidence, and given the rapid accumulation of sequence data for several organisms, exemplars should be avoided when possible, and whole genomes or (at least) annotated TE populations should be preferred for the normalization of M_sRNAs.

#### Impact of normalization on data inference

To gain further insights into how sRNA metrics can change as a function of methodology, we compared the two extremes of a theoretical ‘normalization spectrum’, i.e. un-weighted vs. genome-weighted sRNA data, by investigating two questions of increasing complexity. The first was the general mapping characteristics of TE families, i.e. levels of sRNA targeting per nucleotide of full-length elements. We found that this metric was not altered substantially by the two extremes, because 24nt sRNA targeting remained stronger than 21-22nt sRNAs across most tissues for all families (Figure S3).

The second question was how sRNA mapping correlates with other TE variables. For this question, we studied sRNA targeting as a function of the insertion age of TEs. Age was calculated based on the sequence divergence of each LTR pair and profiled at the family level (Figure 4A). Use of un-weighted data generated strong negative correlations between age and both sRNA species and sRNA expression for all combinations of tissue, family and sRNA length (average Spearman *r* = −0.67, *P* < 10^−20^; Figures 4B, S4). Critically, use of genome-weighted data retained this pattern only for 21-22nt sRNAs (average Spearman *r* = −0.35, *P* < 10^−20^ in most cases), while for 24nt sRNAs there was discordance both between sRNA metrics and among families. We detected a positive correlation for *Ji*, *Opie* and *Huck* using sRNA species, which was often reversed or not statistically supported using sRNA expression (Figures 4B, S4). In contrast, there was a negative correlation for *Cinful-zeon*, *Flip* and *Giepum* across most tissues and for both sRNA metrics.

**Figure 4.**
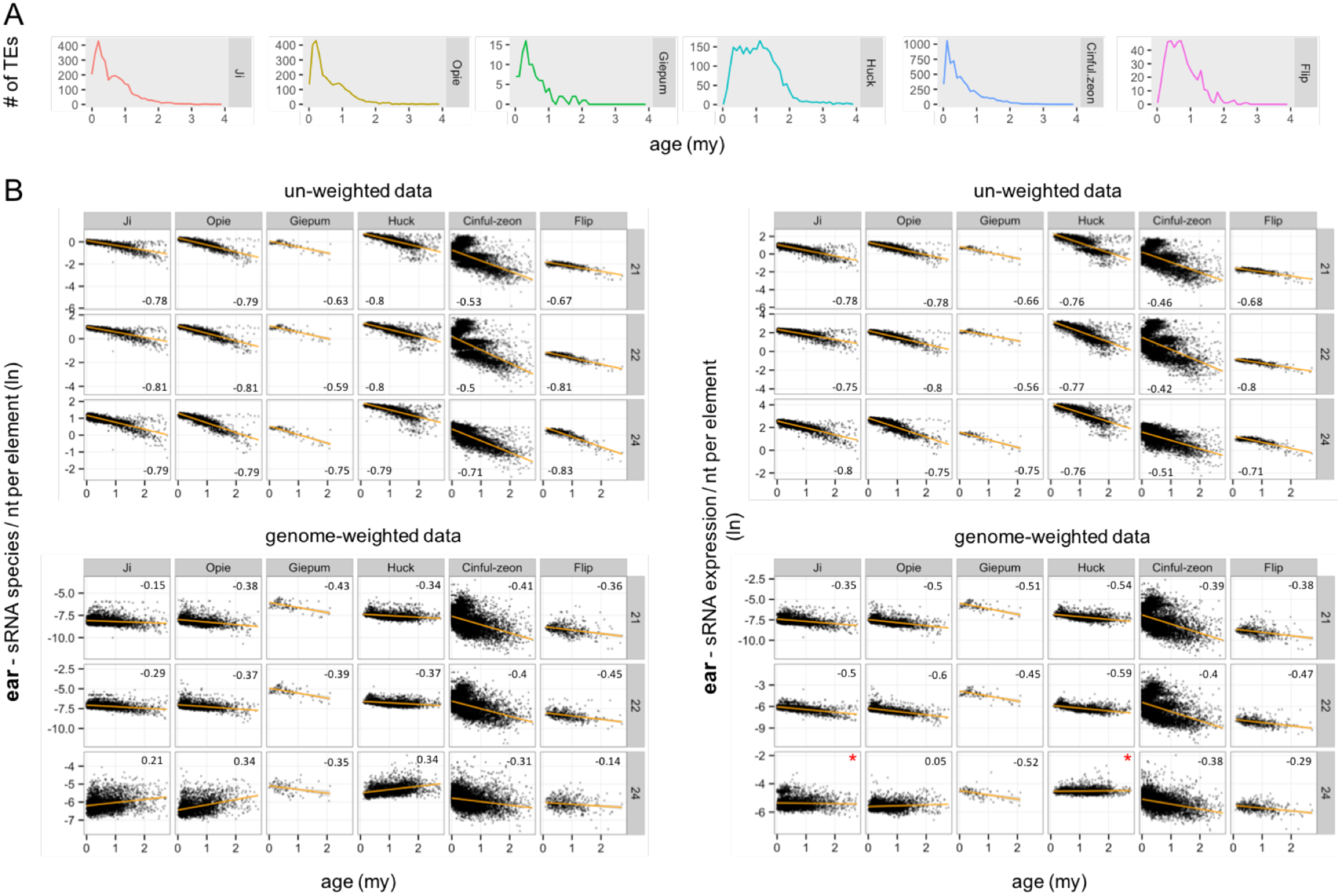
Relationship between TE age and sRNA mapping using un-weighted and genome-weighted approaches. (**A**) Age distribution in million years (my) of TE families. (**B**) Mapping of sRNA species (left panels) or expression (right panels) from ear tissue was calculated per nucleotide of full-length elements for each family. Age is cutoff at 3my to allow sufficient visualization of the *x*-axis. The Spearman *r* coefficient is shown for each plot, calculated for all elements and not only for those <3my. *P* values were <0.01, except those indicated by an asterisk.

Finally, we contrasted the above analysis with how normalization affects another type of commonly used HTS libraries, that is of mRNA expression data. It is known that mRNA libraries have a considerably smaller amount of multiple mapping reads (10% against the genome, in comparison to 40-90% of sRNA libraries), which can be attributed to the longer read lengths and mostly single-copy transcription loci (23). We therefore retrieved mRNA data from three biological leaf replicates and examined (as we did with sRNAs) i) their general mapping characteristics to the genome, ii) the expression patterns of TE families, and iii) the relationship between expression and TE age. First, our analyses confirmed the aforementioned ~10% of M_mRNAs at the genome levels, but we additionally discovered that the vast majority TE-mapping reads were M_mRNAs (Table S4). Furthermore, the median number of locations for these TE-mapping M_mRNAs across the genome or within the annotated full-length elements (Table S4) was approximately only two-fold lower to those of the TE-mapping M_sRNAs (Table 1).

Second, both normalization approaches generated the same relative expression levels among families despite their widely different sizes (Figure S5A). Finally, both types of data produced strong negative correlations between mRNA expression and age for all possible combinations (average Spearman *r* = −0.61, *P* < 10^−20^; Figure S5B).

Taken together, these findings suggest that the choice of treatment of HTS data can affect biological inference, clearly evident in the inconsistent relationship between 24nt sRNAs and age. And although the conclusions of many of the analyses were unchanged, we note that the strength of the correlations with age were substantially weaker for genome-weighted than un-weighted data (average *r* of 0.32 vs. 0.67 for sRNAs and 0.72 vs. 0.52 for mRNAs, using absolute values). This is counterintuitive, because, as we showed earlier (Table 1), weighting-by-location is expected to have a stronger impact on high-copy than low-copy sequences. Yet, 21-22nt sRNA and mRNA profiles did not change as a function of age within each family, whereby the numerous young and highly similar elements were mapped by more sRNAs (Figures 4B, S4) or mRNAs (Figure S5B) than their few, old and divergent relatives in both normalization approaches. Also, the relative mRNA expression levels of families of varying sizes did not change by using weighted or un-weighted data (Figure S5A). Based on these insights and the unpredictability of how normalization may further impact other research questions, we argue that multiple approaches should be used in parallel to validate results.

#### U_sRNA-guided mapping of M_sRNAs may be problematic for TE studies

An alternative approach for mapping M_sRNAs assigns reads to single loci using as guide the local densities of U_sRNAs (23). This method, which is at the core of the ShortStack tool (12), aims to find the true generating locus of each read, instead of allocating M_sRNAs across their targets or even excluding them altogether. Historically, this concept was initially tested with mRNA data where it significantly improved placement of M_mRNAs (48). For sRNAs, recent analysis of simulated libraries by Johnson et al. (2016) showed that the U_sRNA-guided mode outperforms other methodologies in selecting the correct locus from which an M_sRNA may have originated.

However, our data suggest that two properties of TEs may pose a real challenge to this process. First, there is a very small number of U_sRNAs that align to TEs. For example, only 2,166 of 147,034 sRNA species of the ear library that collectively mapped to our *Copia* and *Gypsy* elements are U_sRNAs (Figure 2B, Table S2); furthermore, the vast majority of these U_sRNAs mapped to different TEs (Figure 2D). As a result, and given that the length of our TEs ranges between 7-15kb and that ShortStack examines 250nt windows (23), it is expected that most windows will not have a U_sRNA score and hence vast amounts of M_sRNAs will be discarded. The second issue concerns the numerous genomic locations for M_sRNAs targeting TEs (Table 1). These are far above the 50-target cutoff that Johnson et al. suggest leads to a high rate of misplacement.

Finally, ShortStack can also guide M_sRNA allocation by calculating the densities of both U_sRNAs and weighted M_sRNAs; however, this option did not perform as well as the U_sRNA-only option at the genome level in Arabidopsis, rice and maize (23) and, hence, it is likely that its performance will be further compromised in TE-focused analyses.

A last important point is that a distinction should be made between sRNA-generating vs. sRNA-targeting loci. ShortStack appears to work beautifully for allocating M_sRNAs to their single locus of origin, and future developments may make this approach more efficient for TE data as well. Nonetheless, studies that investigate sRNA targeting patterns may benefit more by methods that allow multiple mapping. This may be especially important for TEs, where it is possible that a given sRNA mediates silencing of more than one locus. Although not empirically proven yet, this conjecture is supported by evidence for the importance of M_sRNAs in RdDM (34), the homology-based *trans* silencing pathway among TEs (38), and the cytoplasmic step of Argonaute loading that dissociates sRNAs from their generating loci (49).

### sRNA metrics: unexpected differences between sRNA species and sRNA expression

So far, our analysis has indicated that sRNA species and sRNA expression generally produce similar results. However, this is not always true. When we examined the relationship between sRNAs and age separately for the LTRs and the internal (INT) domain of TEs using un-weighted data, we observed that the plots of the *Opie* family were markedly different in one case. The expression levels of 24nt sRNAs from leaf on the LTRs split the *Opie* elements in two distinct groups, whereby the ‘upper zone’ was mapped by approximately twice as many reads compared to the ‘lower zone’ (Figure 5A). Species of 24nt sRNAs did not generate the same pattern, nor did other combinations of sRNA lengths and metrics in *Opie* (Figure 5A), or in other families or tissues (not shown).

**Figure 5.**
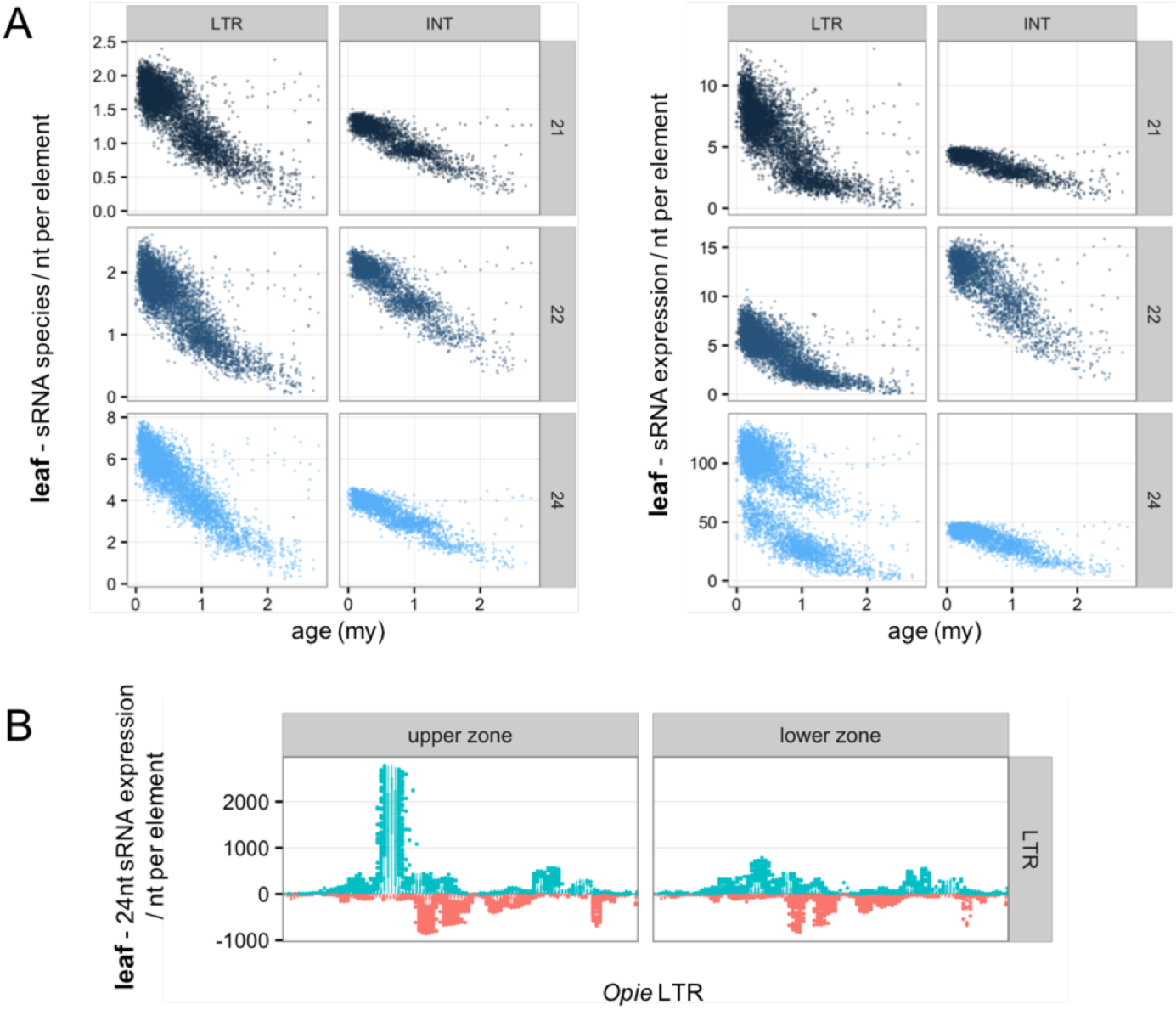
*Opie* population split based on sRNA expression data from leaf tissue. (**A**) Relationship between TE age and number of sRNA species (left) or expression (right) calculated per nucleotide of the *Opie* LTRs and INT domain. Age is cutoff at 3my to allow sufficient visualization of the *x*-axis. (**B**) Mapping patterns (calculated as in Figure 3A) of 24nt expression data along the LTRs of the two distinct *Opie* subpopulations. sRNA data in **A** and **B** were not weighted by their number of genomic loci.

To our surprise, closer investigation revealed that this ‘zoning’ was triggered by sRNAs that mapped to a narrow region on the sense strand of the LTRs (Figure 5B). This region was targeted by ~115x more reads in the elements of the upper zone compared to the elements of the lower zone (median coverage of 1,610 and 14 reads/nt respectively), while there was only a three-fold difference (6.1 vs. 2.1 reads/nt) along the rest of the LTR. This finding implies that highly expressed sRNA species mapping to this region of the elements of the upper zone may drive the *Opie* split. We retrieved 836 24nt sRNA species from all *Opie* elements and, surprisingly, only one appeared to be responsible for the zoning. This sRNA species combined very high expression (1,976 reads) and number of target LTRs (3,228 LTRs that comprise the upper zone), ranked 1^st^ and 7^th^ respectively among the 836 sRNAs. In contrast, most other highly-mapped sRNAs of the same region had expression levels of <10 reads, and therefore did not contribute to the zoning.

The fact that a single sRNA generates this spectacular pattern raises several methodological concerns. First, it is likely that such very high expression levels may be the outcome of biases during library construction (15). Second, our data imply that the use of sRNA species is more robust than sRNA expression, because it appears to be less sensitive to errors that can occur, e.g., during PCR amplification. Finally, and perhaps most importantly, these findings denote the need for the confirmation of such observations. This can be achieved by cross-examining results from different normalization approaches. We checked if genome-weighted sRNA expression data reproduce the zoning, but this does not seem to be the case (not shown). However, given the inconsistencies of normalization approaches as discussed previously, the most appropriate way is the inclusion in the experimental design of technical and/or biological replicates. In previous years the lack of sRNA studies could be attributed to the high costs of sequencing. These costs are now much lower and, hence, replicates should be typically included in epigenetic studies to help identify aberrancies.

## CONCLUDING REMARKS

In this work we attempted to address the complex issue of mapping and analyzing sRNAs in the context of TEs, which comprise the vast majority of most plant genomes. With the goal of providing insights that might help guide future studies, we compared mapping to TE exemplars vs. annotated TE populations; explored the extent of multiple mapping across different genomic regions or datasets; examined how various strategies for mapping M_sRNAs (or M_mRNAs) affect biological inference or are applicable to TEs; and finally presented an unexpected inconsistency between the two sRNA metrics of species and expression. Summarizing each section into take-home messages:

1. TE exemplars have been widely popular thus far for various reasons, including the absence of sufficient sequence information or, indeed, because research would not truly benefit from the burdensome analysis of annotated TE populations. However, their usage comes with several limitations that scientists need to be aware of, such as the presence of unresolved/masked regions in their sequences, the overrepresentation of U_sRNAs, and the large volumes of unmapped data that may severely bias analyses.
2. Annotated TE populations appear to be more informative than exemplars for mapping epigenetic data. Given that sequence data are now available for a large number of genomes, the use of TE populations – additionally filtered for foreign DNA if needed – should be preferred over exemplars when possible.
3. Our analyses also suggest that the inclusion of M_sRNAs in TE studies is necessary, and we strongly advocate against a focus solely on U_sRNAs. We favor using comparative un-weighted and weighted (i.e. normalized) mapping approaches in parallel to validate biological inferences. Crucially, whole, or even partially, sequenced genomes should be preferred over exemplars for weighting M_sRNAs. Furthermore, approaches that assign M_sRNAs to single loci based on U_sRNAs density are very promising but are not yet applicable for TE studies.
4. Finally, the metric of sRNA expression (and to a lesser extent sRNA species) may be prone to errors during HTS library construction. However, sRNA expression is often a crucial measurement, for example during differential expression analysis. Aided by the rapidly falling costs of sequencing, the inclusion of technical and/or biological replicates in sRNA studies should now be standard.

## METHODS

### TE datasets

For the *Ji*, *Opie* and *Giepum* Sirevirus families of the *Copia* superfamily we used the strictly curated set of full-length elements that were previously analyzed in Bousios et al. 2016. We also retrieved the complete annotated populations from the MASiVEdb website (http://bat.infspire.org/databases/masivedb/) (31).

For the *Huck*, *Cinful-zeon* and *Flip* families of the *Gypsy* superfamily we first retrieved the repeat annotation file of the maize TE consortium (ZmB73_5a_MTEC+LTR_repeats.gff) from ftp.gramene.org. This file, however, does not specify whether an annotated region represents full-length or fragmented TEs. Hence, we plotted the frequency distribution of the lengths of the annotated regions to identify peaks for each family that would correspond to the size of full-length elements as calculated by Baucom et al. (32) (Figure S6A). This approach identified a single peak for *Huck* that nearly overlapped with the Baucom full-length average (13.4kb), two peaks for *Cinful-zeon* that flanked the Baucom average (8.2kb), and two peaks for *Flip* – one nearly overlapping with the Baucom average (14.8kb) and one residing in close proximity (Figure S6A). Based on these results, we selected regions between 13.3-14.1kb for *Huck*, 7.1-7.5kb and 9.2-9.7kb for *Cinful-Zeon*, and 14.8-15.6kb for *Flip* as candidates for full-length elements, retrieving 2,614, 6,965 and 607 sequences respectively. We then ran LTRharvest (50) with parameters *xdrop* 25, *mindistltr* 2000, *maxdistltr* 20000, *ins* −3, *del* −3, *similar* 50, *motif* TGCA, *motifmis* 1, *minlenltr* 100, and *maxlenltr* 5000 in order to identify the borders between the LTRs and the INT domain, and to also calculate the canonical LTR length of each family. Based on our approach, we selected LTR lengths between 1-1.8kb for *Huck*, 450-750nt for *Cinful-zeon*, and 4.1-4.5kb for *Flip*, finally yielding 2,460, 6,276 and 483 full-length elements for each family respectively (Figure S6B).

The insertion age of each *Copia* and *Gypsy* TE was calculated by first aligning the LTRs using MAFFT with default parameters (51) and then applying the LTR retrotransposon age formula with a substitution rate of 1.3 x 10-8 mutations per site per year (52).

Finally, all maize TE exemplars were downloaded from http://maizetedb.org. Note that we removed one *Ji* (RLC_ji_AC186528-1508) and two *Giepum* (RLC_giepum_AC197531-5634; RLC_giepum_AC211155-11010) exemplars from our analysis, based on evidence from Bousios et al. (2012) that indicated that the specific exemplars are not true representatives of these families.

### Mapping sRNA and mRNA libraries

We used published sRNA data from leaf (GSM1342517), tassel (GSM448857), and ear (GSM306487) tissue, and mRNA data from three technical replicates from leaf (SRR531869, SRR531870, SRR531871) tissue. Adapters and low quality nucleotides were removed using Trimmomatic and the FASTX toolkit respectively, until every read had three or more consecutive nucleotides with a Phred quality score of >20 at the 3’-end. The libraries were filtered for miRNAs (http://www.mirbase.org/), tRNAs (http://gtrnadb.ucsc.edu/), and rRNAs and snoRNAs (http://rfam.sanger.ac.uk/); sRNA reads of 21nt, 22nt and 24nt length and mRNA reads longer than 25nt were mapped to the maize B73 genome (RefGen_V2) and the maize TE database using BWA with default settings and no mismatches (22). Following previous work (26), each distinct sRNA or mRNA sequence was termed “species”, and the number of its reads was its “expression”.

After mapping, each species was tagged as either U_sRNA/U_mRNA or M_sRNA/M_mRNA separately for the genome and the exemplar database. M_sRNAs/M_mRNAs were either normalized by their number of mapping locations or not normalized, depending on the analysis. Finally, we calculated the total number of sRNA species that mapped to a TE ‘locus’ (i.e. the full-length sequence, LTRs or INT domain), but also the number of sRNA species and sRNA expression (weighted or un-weighted) per nucleotide of each locus. The per nucleotide measures allow comparisons of averages among TEs and also analysis along the length of the TE sequence.

## SUPPLEMENTAL MATERIAL

The sequences and other information of the annotated TE populations are available as Supplemental Material in bat.infspire.org/sireviruses/Submission_Data/.

## ACKNOWLEDGMENTS

AB is supported by the European Community’s Seventh Framework Programme (FP7/2007-2013) under grant agreement no: PIEF-GA-2012-329033; BSG by NSF grant IOS-1542703 and a fellowship from the Borchard Foundation; ND by research grant AZV-MZ-CR 16-34272A-4/2016 and project CEITEC 2020 (LQ1601).

